# Muscle effort is best minimized by the right-dominant arm in the gravity field

**DOI:** 10.1101/2021.07.13.452146

**Authors:** Gabriel Poirier, Mélanie Lebigre, France Mourey, Charalambos Papaxanthis, Jeremie Gaveau

## Abstract

The central nervous system (CNS) is thought to develop motor strategies that minimize various hidden criteria, such as end-point variance or effort. A large body of literature suggests that the dominant arm is specialized for such open-loop optimization-like processes whilst the non-dominant arm is specialized for closed-loop control. Building on recent results suggesting that the brain plans arm movements that takes advantage of gravity effects to minimize muscle effort, the present study tests the hypothesized superiority of the dominant arm motor system for effort minimization. Thirty participants (22.5 ± 2.1 years old; all right-handed) performed vertical arm movements between two targets (40° amplitude), in two directions (upwards and downwards) with their two arms (dominant and non-dominant). We recorded the arm kinematics and the electromyographic activity of the anterior and posterior deltoid to compare two motor signatures of the gravity-related optimization process; i.e., directional asymmetries and negative epochs on phasic muscular activity. We found that these motor signatures were still present during movements performed with the non-dominant arm, indicating that the effort-minimization process also occurs for the non-dominant motor system. However, these markers were reduced compared with movements performed with the dominant arm. This difference was especially prominent during downward movements, where the optimization of gravity effects occurs early in the movement. Assuming that the dominant arm is optimal to minimize muscle effort, as suggested by previous studies, the present results support the hypothesized superiority of the dominant arm motor system for effort-minimization.

## Introduction

Handedness is a particular aspect of motor control referring to the preferential employment of a hand – the dominant one – to perform daily-life tasks. Asymmetries in the neural organization of cerebral hemispheres could explain inter-limb differences in the control of arm movements (Carson et al. 1992; Flowers 1975; Jayasinghe et al. 2021; Roy et al. 1994; Roy and Elliott 1986; Sainburg 2002, 2014; Woytowicz et al. 2018). A large body of literature suggests that the dominant arm/hemisphere is specialized for open-loop optimization-like processes, whilst the non-dominant arm/hemisphere is specialized for closed-loop control that stabilized arm postures against unexpected environmental conditions (Bagesteiro and Sainburg 2002; Sainburg 2002; Schaffer and Sainburg 2017; Yadav and Sainburg 2011, 2014). For example, when asking participants to perform reaching movements in predictable and unpredictable force fields, Yadav and Sainburg (2014) observed that the dominant arm performed better in the predictable force field whereas the non-dominant arm outperformed the dominant one in the unpredictable force field.

Hitherto, most studies assessed dominant vs non-dominant arm control comparing movements performed into a transverse plane, with the arm’s weight supported by the experimental device (i.e., compensating for gravity mechanical effects). Recently, Schaffer and Sainburg (2017) investigated unsupported horizontal arm movements, making arm excursions along the vertical axis possible but redundant to the task. The authors found that vertical excursions were greater for the dominant than the non-dominant arm, and correlated with movement errors for the non-dominant but not the dominant arm. The authors proposed that these results reflect different control strategies in which only the dominant arm/hemisphere can take advantage of gravity effects during the movement.

Investigating motor planning and control of the dominant arm/hemisphere, several studies proposed that the brain uses an internal representation of gravity to take advantage of its mechanical effects (Berret et al. 2008; Crevecoeur et al. 2009; Gaveau et al. 2014, 2016, 2021; Gaveau and Papaxanthis 2011). More specifically, studies investigating vertical arm reaching movements reported direction-dependent arm kinematics (Gaveau et al. 2011, 2014, 2016, 2021; Gaveau and Papaxanthis 2011; Gentili et al. 2007; Hondzinski et al. 2016; Papaxanthis et al. 2005; Poirier et al. 2020; Le Seac’h and McIntyre 2007; Yamamoto et al. 2016, 2019; Yamamoto and Kushiro 2014). Particularly, the time to peak acceleration and time to peak velocity were shorter and the curvature was greater for upward than for downward movements. These directional differences progressively disappeared during microgravity exposure (Gaveau et al. 2016; Papaxanthis et al. 2005) and well-matched the results of simulation models that optimize gravity effects to minimize muscle effort (Berret et al. 2008; Crevecoeur et al. 2009; Gaveau et al. 2014, 2016). This optimization process was recently further supported by the observation of specific negative epochs on the phasic activity of antigravity muscles (Gaveau et al. 2021). Direction-dependent kinematics and negative epochs on the phasic activity of muscles are thought to represent the hallmark of an optimization process that takes advantage of gravity torque to minimize muscle effort during vertical arm movements.

Building on the motor control literature on arm lateralization and on gravity-related muscle effort minimization, the present study tested the hypothesized superiority of the dominant arm motor system – compared to the non-dominant one – for gravity-related muscle effort minimization. We compared kinematic and electromyographic patterns of vertical arm reaching movements performed with the dominant and the non-dominant arm. We expected motor signatures of the gravity-related optimization process (i.e., directional asymmetries and negative phases in phasic EMG) to be larger for the dominant than for the non-dominant arm.

## Materials and Methods

### Participants

Thirty healthy young volunteers [17 males and 13 women; mean age = 22. 5 ± 2.1 (SD) years; mean height = 173 ± 8 cm; mean weight = 70 ± 10 kg] were included in this study after giving their written informed consent. Participants had normal or corrected-to-normal vision and did not present any neurological or muscular disorders. The laterality index of each participant was superior to 60 (Edinburgh Handedness Inventory, Oldfield 1971), indicating that all participants were right-handed. The study was carried out following legal requirements and international norms (Declaration of Helsinki, 1964) and approved by the French National ethics committee (2019-A01558-49).

### Experimental protocol

Our experimental protocol was similar to those of previous studies investigating gravity-related arm movement control (Gaveau et al. 2014, 2016; Gaveau and Papaxanthis 2011; Gentili et al. 2007; Poirier et al. 2020; Le Seac’h and McIntyre 2007). Participants were asked to perform unilateral single-degree-of-freedom vertical arm movements (rotation around the shoulder joint) in a parasagittal plane with their dominant (D) and non-dominant (ND) arms. We investigated single-degree-of-freedom vertical arm movements to isolate and emphasize the mechanical effects of gravity on arm motion. During such movements, the gravity torque varies with movement direction, whereas inertia remains constant. Movements with the D and ND arms were organized in a random block design. Upward and downward directions were randomly interleaved within each block. Each participant carried out 120 trials (30 trials x 2 directions x 2 arms).

Participants sat on a chair with their trunk vertically aligned (Fig.1A). Two targets (diameter of 3 cm) were placed in front of the participant’s right or left shoulder (parasagittal plane), at a distance corresponding to the length of their fully extended arm plus two centimeters. The required angular shoulder rotation between the two targets was 40° (Fig.1A), corresponding to a 110° (upward target) and 70° (downward target) shoulder elevation.

**Figure 1.**
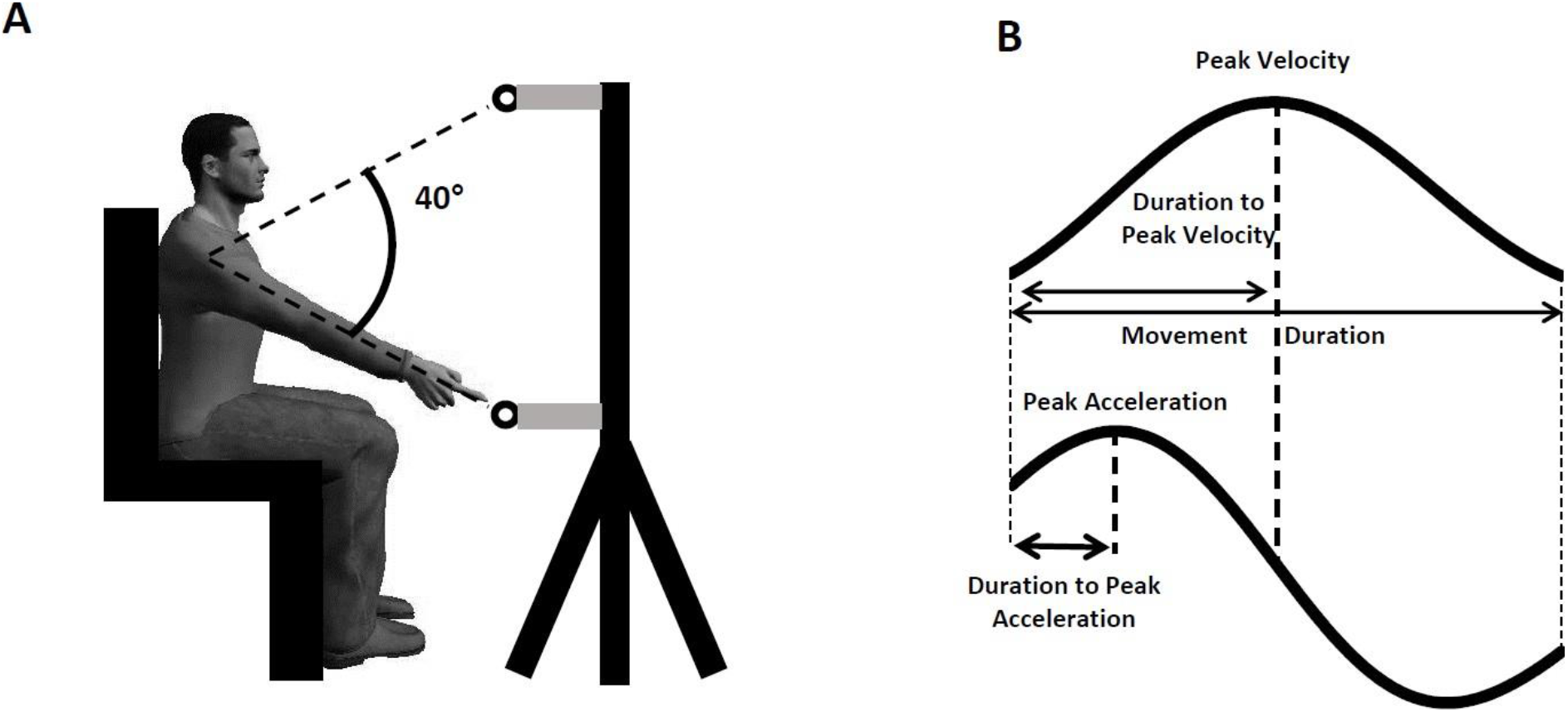
(A) Experimental setup. Participants performed arm pointing movements between two targets on separated trials. (B) Illustration of the parameters computed on the velocity and acceleration profiles.

Each trial was carried out as follows: the experimenter indicated the starting target (red for upward or blue for downward) and the participant positioned his arm fully extended in front of it (initial position). After a brief delay (~2 seconds), the experimenter verbally informed the participant that she/he was free to aim at the other target, as fast and accurately as possible, whenever she/he wanted. Note that reaction time was not emphasized in our experiment. Participants were requested to maintain their final position for a brief period (about 2 seconds) until the experimenter instructed them to relax their arm near the hips. A short rest period (~10 s) separated trials to prevent muscle fatigue. Additionally, a 5mn time interval separated each block. Participants were allowed to perform few practice trials (~5 trials) before each block.

Five reflective markers were placed on the participant’s shoulder (acromion), arm (middle of the humeral bone), elbow (lateral epicondyle), wrist (cubitus styloid process), and finger (nail of the index). We also placed a marker on each target to measure end-point error. Position was recorded with an optoelectronic motion capture system (Vicon system, Oxford, UK; six cameras) at a sampling frequency of 100Hz. The spatial precision of the system was less than 0.5mm. Additionally, we placed two bipolar surface electrodes (Aurion, Zerowire EMG, sampling frequency: 1000Hz) on the anterior (DA) and posterior (DP) heads of the deltoid to record EMG activity. The Giganet unit (Vicon, Oxford, UK) recorded kinematic and EMG signals synchronously.

### Data analysis

We processed Kinematic and EMG data using custom programs written in Matlab (Mathworks, Natick, NA). Data processing was similar to previous studies (Gaveau et al. 2021; Poirier et al. 2020).

#### Kinematics analysis

First, we filtered position using a third-order low-pass Butterworth filter (5Hz cut-off, zero-phase distortion, “butter” and “filtfilt” functions). We then computed velocity and acceleration profiles by deriving position signals. Angular joints displacements were computed to ensure that rotations, apart from the shoulder joint, were negligible (<1° for each trial). We defined movement onset and offset as the moments when finger velocity rose above or fall below, respectively, a threshold corresponding to 10% of peak velocity. Movements presenting multiple local maxima on the velocity profile were automatically rejected from further analysis (< 1% of all trials). The following parameters were then computed from angular profiles of the virtual segment linking finger to shoulder markers (Fig.1B): 1) Movement duration (MD), defined as the duration between movement onset and offset. 2) Movement amplitude (Amp), defined as the angular amplitude between movement onset and offset. 3) Relative duration to peak acceleration (rD-PA), defined as the time to peak acceleration normalized by total movement duration. 4) Relative duration to peak velocity (rD-PV), defined as the time to peak velocity normalized by total movement duration. 5) Distance between finger end-point position and target position (spatial error). Because the task was to perform vertical single degree-of-freedom movements, this distance was calculated on the vertical axis only. For each participant and condition, constant and variable errors were calculated as follows:

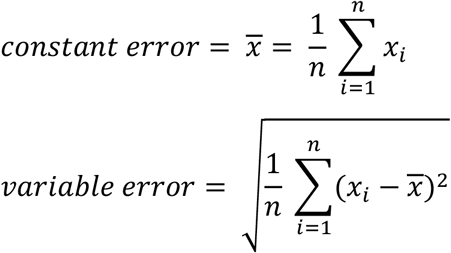

Where n is the total number of trials in the condition (30 for each condition), *x_i_*, is the spatial error between finger end-point position and target position for the i^th^ movement of the condition, and 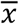 is the constant error in the condition.

#### EMG analysis

EMG signals were first rectified and filtered using a bandpass third-order Butterworth filter (bandpass 20-300Hz, zero-phase distortion, “butter” and “filtfilt” functions). Signals were integrated using a 50ms sliding window and cut off from 250ms before movement onset to 250ms after movement offset. Then, signals were filtered using a low-pass third-order Butterworth filter (low-pass frequency: 20Hz) and normalized by the maximal EMG value recorded during maximal voluntary isometric contractions (MVIC). At the beginning of each block, participants performed three flexions and three extensions MVIC at a shoulder angle of 90°. Lastly, EMG traces were classified according to movement mean velocity and averaged across three trials (from the three slowest to the three fastest movements), resulting in 10 EMG traces to be analyzed for each block. Each set of three traces was normalized in duration (corresponding to the mean duration of the three traces) before averaging.

We computed the phasic component of each EMG signal using a well-known subtraction procedure (Buneo et al. 1994; D’Avella et al. 2006, 2008; Flanders et al. 1994; Flanders and Herrmann 1992; Gaveau et al. 2021). We averaged the values of the integrated EMG signals from 500ms to 250ms before movement onset and from 500ms to 250ms after movement offset (Fig. 2A). The tonic component was calculated with a linear interpolation between these two averaged values (Fig.2B). Finally, we computed the phasic component by subtracting the tonic component from the full EMG signal (Fig.2C).

**Figure 2.**
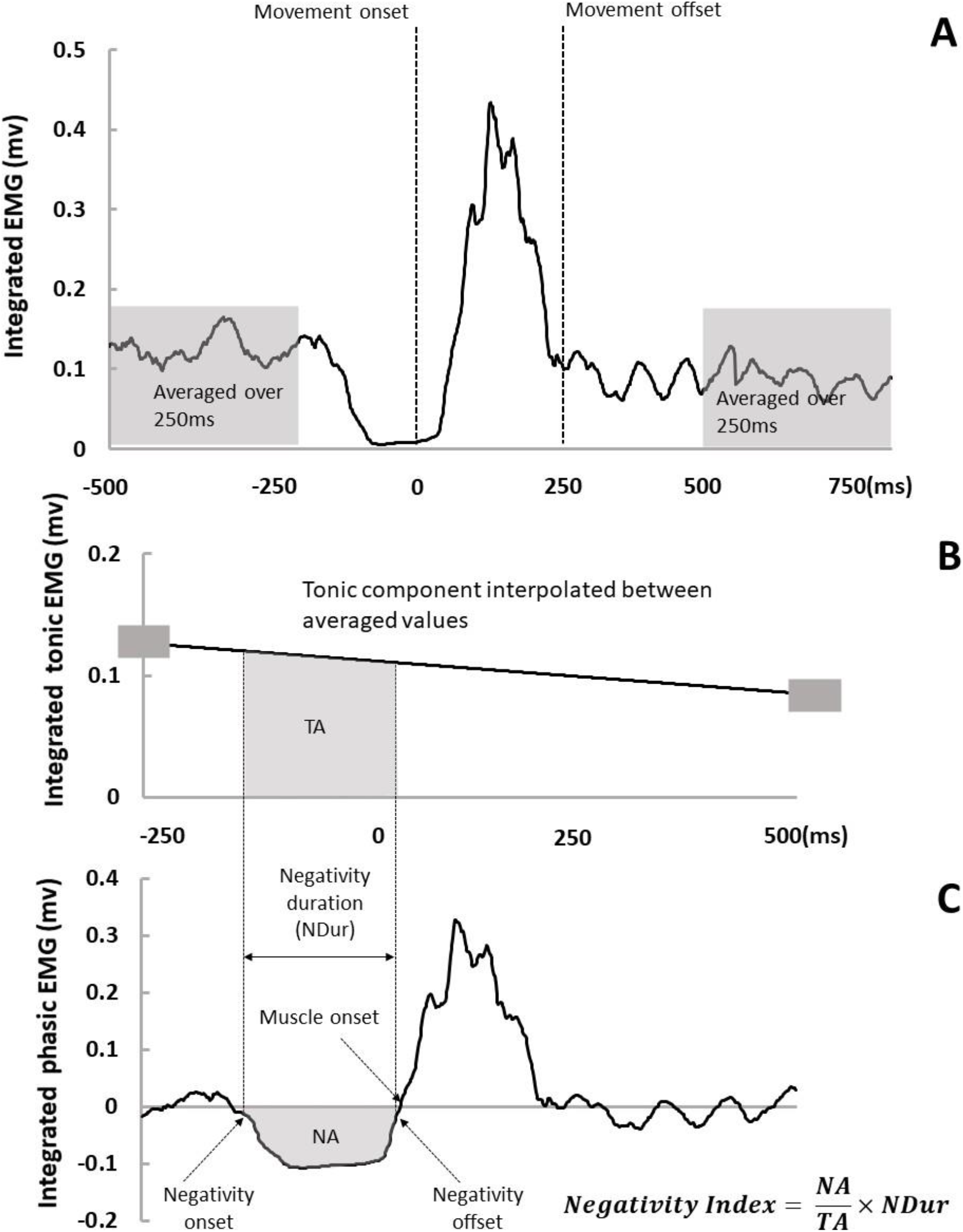
Analysis of the tonic and phasic components of the EMG. (A). Illustration of the integrated full EMG signal. Movement onset and offset were detected on the velocity profile. We averaged the signal from 500 to 250ms before movement onset and from 250 to 500ms after movement offset (grey windows). (B). Tonic EMG component was obtained by linear interpolation between the two previously averaged values (small grey rectangles). (C). Phasic EMG component was obtained by subtracting the tonic component (in panel B) from the full EMG (in panel A). We detected negativity onset and offset and then computed negativity duration (NDur) and integrated phasic signal between negativity onset and offset (NA). TA in panel B represents the integrated tonic signal in the same period.

Using a 95% confidence interval threshold on the baseline values (signal from 500 to 250ms before movement onset), we defined the onset of phasic muscle activation bursts and computed the following parameters on phasic EMG signals (see Fig.2C): 1) Agonist onset, defined as the time of the agonist muscle onset relative to movement onset (in milliseconds), 2) Antagonist onset, defined as the time of the antagonist muscle onset relative to movement onset (in milliseconds), AD was considered as agonist muscle for upward movements and as antagonist muscle for downward movements. Inversely, PD was considered as agonist muscle for downward movements and as antagonist muscle for upward movements. These parameters allow us to compare muscular organization and activity between arms and directions.

It was recently shown that phasic EMG activity of antigravity muscles consistently exhibits negative epochs during vertical arm movements (Gaveau et al. 2021) when gravity is coherent with the arm acceleration sign (in the acceleration phase of downward movement and the deceleration phase of upward movements). This observation likely reflects an optimal predictive motor strategy where muscle activity is decreased when gravity assists arm movements, thereby discounting muscle effort. We defined negative epochs as an interval where the phasic EMG signal was inferior to zero minus the 95% confidence interval (computed on the integrated EMG signals from 500ms to 250ms before movement onset) for at least 50ms. We used this value as a threshold to define negativity onset and offset. We also computed negativity occurrence, defined as the number of trials including a negative period divided by the total number of trials in the condition. We calculated a negativity index to compare the amount of negativity in the phasic EMG signal between conditions (Fig.2C):

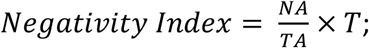

where NA is the integrated phasic signal between negativity onset and offset; TA is the integrated tonic signal between negativity onset and offset (if NA and TA are equal, it means that the muscle is completely silent during the interval); and T is the duration of the negative epoch normalized by movement duration. We also computed negativity duration, defined as the duration of the negative period normalized by movement duration, and negativity amplitude, defined as the minimal 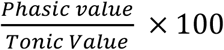 value during the negative period. For example, a value of −100 indicates that the muscle is completely relaxed.

### Statistics

Statistical analyses were performed with STATISTICA 13 (Statsoft Inc., Tulsa, OK, USA). We verified that all variables were normally distributed (Kolgomorov-Smirnov test) and that the sphericity assumption was not violated (Mauchly’s test). We performed repeated measure ANOVA with two factors: *arm* (D vs. ND) and *direction* (Upward vs. Downward). We used *HSD-Tukey tests* for post-hoc comparisons. To specifically compare directional differences between arms when appropriate, we used pre-planned bilateral Student T-tests. The level of significance for all analyses was fixed at *p* = 0.05.

## Results

### Kinematics

All participants performed single-degree-of-freedom vertical arm movements in the parasagittal plane (shoulder azimuth angle <1° for all trials; n = 3600) with bell-shaped, single-peaked velocity profiles, and double-peaked acceleration profiles. Fig.3 provides a qualitative illustration of velocity and acceleration profiles for each arm and direction from a typical participant. Table 1 shows the mean (±SE) values of all kinematic parameters.

**Figure 3.**
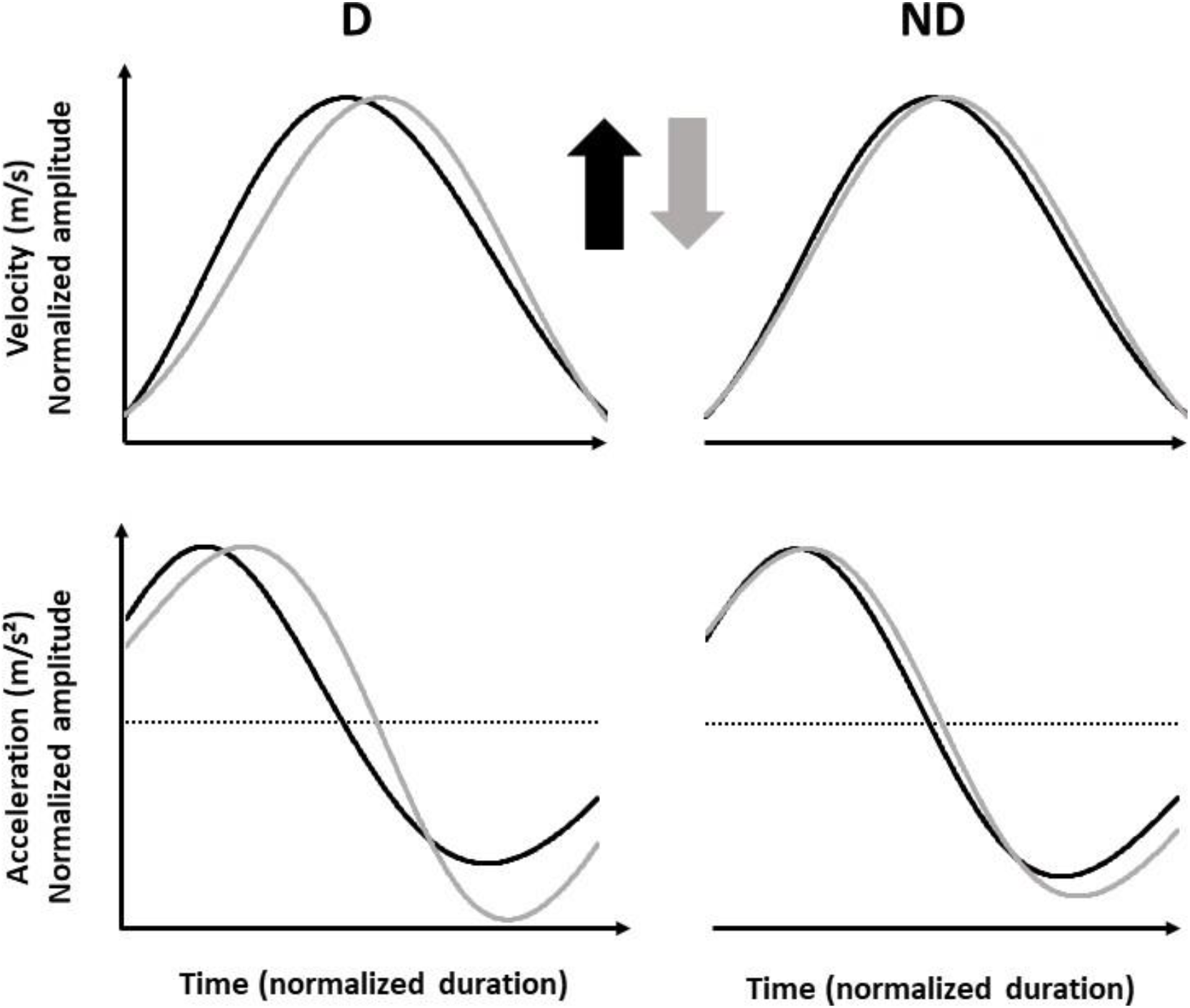
Mean velocity and acceleration profiles for all experimental conditions in a typical participant. Vertical arrows indicate movement directions.

Average angular amplitude was 40.4 ± 3.3° overall. Amplitude varied significantly with *direction* (F = 11.58; p = 2e-3; η_p_^2^ = 0.29), with participants exhibiting slightly higher amplitudes for upward than for downward movements (see movement amplitude in Table.1) but was not affected by *arm* (F = 2.03; p = 0.16; η_p_^2^ = 0.07) or *arm x direction* (F = 0.11; p = 0.08; η_p_^2^ = 0.08). As required by task demands, participants performed fast movements (see movement duration in Table. 1). Neither *arm* (F = 0.19; p = 0.66; η_p_^2^ = 0.007) and *direction* (F = 1.79; p = 0.19; η_p_^2^ = 0.06), nor their interaction affected movement duration (F = 0.07; p = 0.40; η_p_^2^ = 0.02). Thus, similar movement durations and amplitudes with the D and the ND arm ensure that movement speed does not biases the conclusions reached by the following analyses (Gaveau and Papaxanthis 2011).

#### Relative Duration to Peak Acceleration (rD-PA)

Fig.4A displays the mean (+SE) relative duration to peak acceleration (rD-PA) for each arm and direction. *Direction* (F = 46.48; p < 1e-6; η_p_^2^ = 0.62), but not *arm* (F = 1.49; p = 0.23; η_p_^2^ = 0.05), significantly influenced rD-PA. Importantly, we observed a significant *arm x direction* interaction effect (F = 15.05; p = 5.5e-4; η_p_^2^ = 0.34). Post-hoc comparisons highlighted that rD-PA was smaller for upward than downward movement for both the D (p = 1.6e-4; Cohen’s d = 1.26) and the ND (p = 1.6e-4; Cohen’s d = 0.89) arm. It was further revealed that rD-PA was longer with the D than with the ND arm for downward movements (p = 0.003; Cohen’s d = 0.65) but similar between arms for upward movements (p = 0.39; Cohen’s d = 0.29). The *arm x direction* interaction effect demonstrates that the directional asymmetry (the difference between upward and downward) – that has been proposed to represent the hallmark of gravity effects optimization to save muscle effort (Gaveau et al. 2014, 2016, 2021) – differs between the dominant (D) and the non-dominant arm (ND). To better illustrate this effect, we computed a directional difference between directions (Down – Up) for the D and ND arm, separately (see Fig. 4B). A single preplanned bilateral paired t-test confirmed that the directional difference was higher with the D than the ND arm (t = 3.85; p = 5.5e-4; Cohen’s d = 0.64).

**Figure 4.**
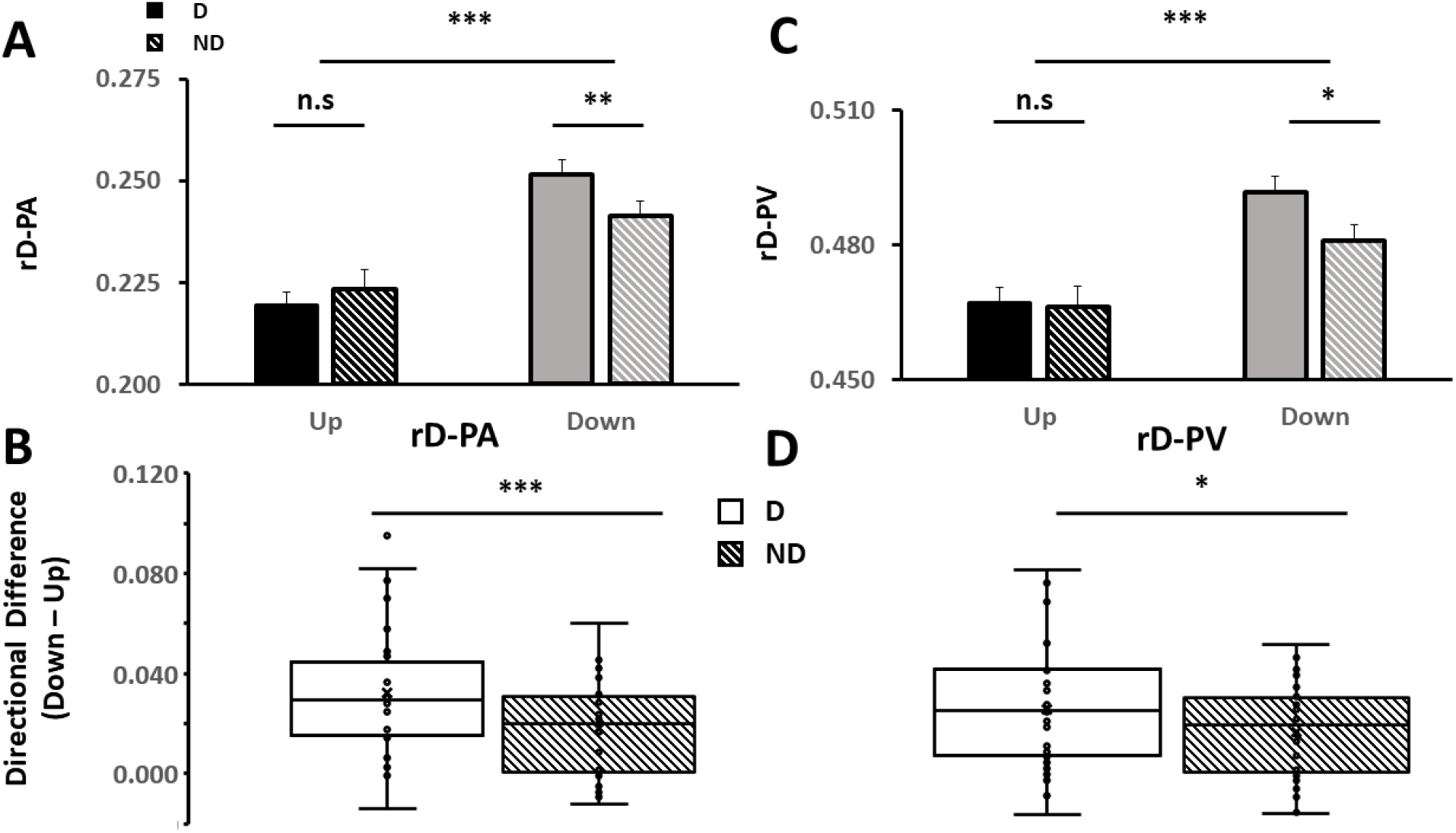
(A). Mean (+SE) relative duration to peak acceleration (rD-PA) for upward (black) and downward (grey) movements. D and ND conditions are represented by respectively solidand striped bars. (B). The box plot shows directional difference on rD-PA (computed for each participant as: rD-PA_Down_-rD-PA_Up_) in the D (white box) and ND (striped box) condition. Each dot represents a participant, the cross represents the mean, and whiskers represent a 95% confidence interval. (C). Mean (+SE) relative duration to peak acceleration (rD-PV) for upward (black) and downward (grey) movements. D and ND conditions are represented by respectively solid and striped bars. (D). The box plot shows directional difference on rD-PV (computed for each participant as: rD-PV_Down_-rD-PV_Up_) in the D (white box) and ND (striped box) condition. Each dot represents a participant, the cross represents the mean, and whiskers represent a 95% confidence interval.

#### Relative Duration to Peak Velocity (rD-PV)

Fig.4C displays mean (+SE) relative duration to peak velocity (rD-PV) for each arm and direction. *Direction* (F = 31.27; p= 5e-6; η_p_^2^ = 0.52) and *arm* (F = 4.44; p = 0.04; η_p_^2^ = 0.13) significantly influenced rD-PV. *Arm x direction* interaction effect (F = 4.74; p = 0.04; η_p_^2^ = 0.14) was also significant, showing once again that directional asymmetries were larger with the D than with the ND arm. Post-hoc analysis displayed that rD-PV was higher for downward than for upward movements in D (p = 1.6e-4; Cohen’s d = 0.99) and ND (p = 5e-4; Cohen’s d = 0.76) conditions. It was also revealed that rD-PV differed between arms in downward (p = 0.01; Cohen’s d = 0.59) but not in upward (p = 0.99; Cohen’s d = 0.05) movements (see Fig.4C). Fig.4D displays the directional difference for both arms. Directional difference was significantly larger with the D than the ND arm (t = 2.17; p = 0.04; Cohen’s d = 0.42).

#### Constant error

Mean (+SE) constant errors for both arms and directions are displayed in Fig.5A. *Direction* significantly influenced end-point constant error (*direction* effect: F = 36.68; p = 2e-6; η_p_^2^ = 0.57). Participants overshot the target during upward movements (resulting in a constant error >0) whereas they undershot the target during downward movements (resulting in a constant error <0). However, neither *arm* (F = 0.28; p = 0.60; η_p_^2^ = 0.01) nor *arm x direction* (F = 0.15; p = 0.70; η_p_^2^ = 0.005) effects were significant.

**Figure 5.**
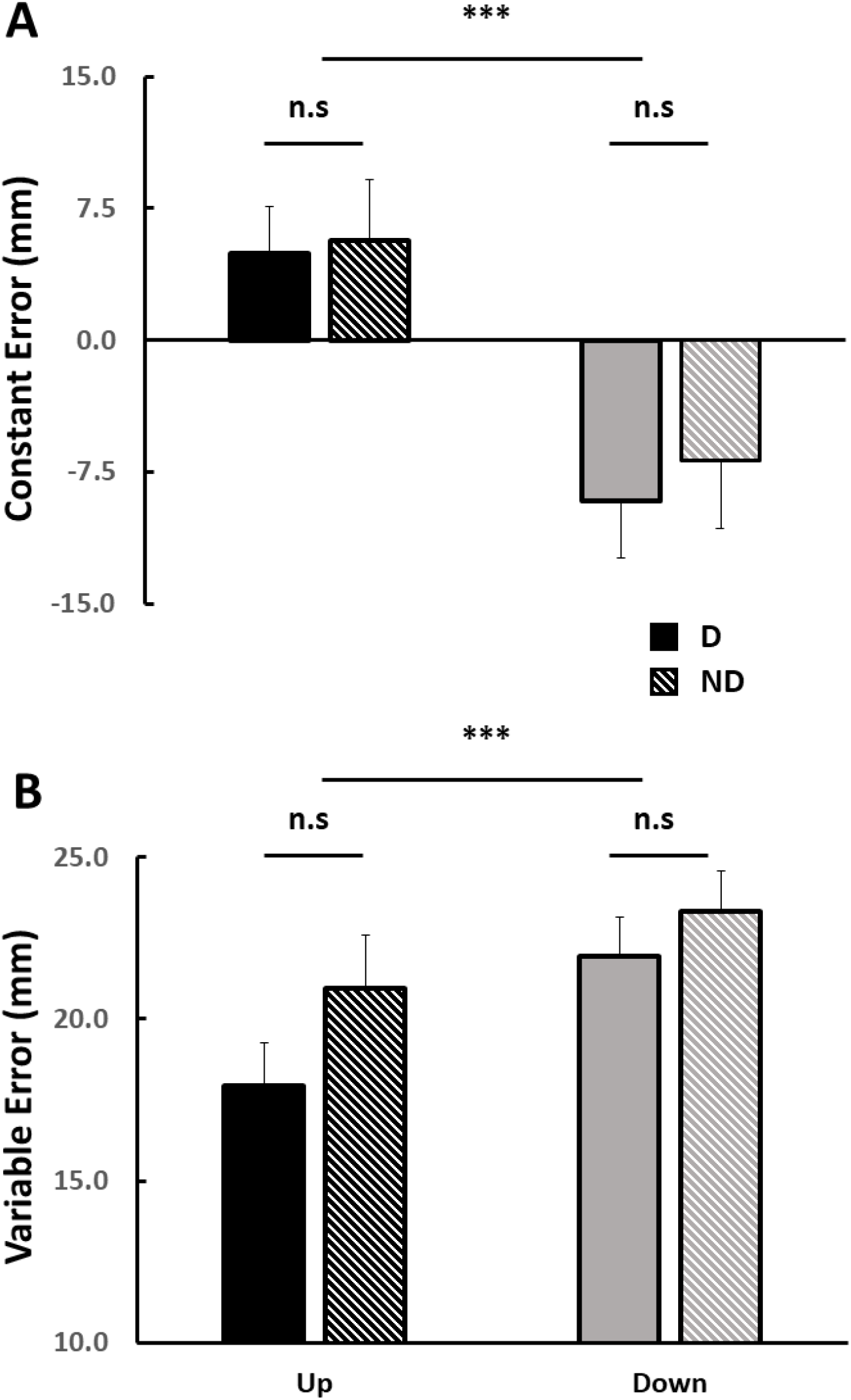
(A). Mean (+SE) constant error for upward (black) and downward (grey) movements. D and ND conditions are represented by respectively solid and striped bars. (B). Mean (+SE) variable error for upward (black) and downward (grey) movements. D and ND conditions are represented by solid and striped bars, respectively.

#### Variable error

Mean (+SE) variable errors for both arms and directions are displayed in Fig.5B. We observed a significant *direction* effect (F = 24.97; p = 2.6e-5; η_p_^2^ = 0.46) revealing that variable error was more important for downward than for upward movements. Neither *arm* (F = 2.42; p = 0.13; η_p_^2^ = 0.08) nor *arm x direction* (F = 1.85; p = 0.18; η_p_^2^ = 0.06) effects were significant.

Overall, kinematic analyses reveal that directional differences in the temporal organization (rD-PA & rD_PV) of vertical arm movements are less important with the non-dominant than with the dominant arm. These results suggest that motor planning leads to less efficient motion with the ND than with the D arm in the gravity field.

### EMG

Fig.6 displays the mean phasic EMG profiles for each muscle, direction, and arm of a typical participant. As recently reported (Gaveau et al. 2021), phasic EMG signals present negative phases during the deceleration of an upward movement and the acceleration of a downward movement; i.e., when gravity torque helps to produce the arm’s motion. This negativity means that the muscle is less activated than it should have been to exactly compensate for gravity torque. Although observable on gravity-muscles too – revealing a strong inactivation of all muscles during these specific phases – negativity is especially prominent in the antigravity-muscle (Anterior deltoid).

**Figure 6.**
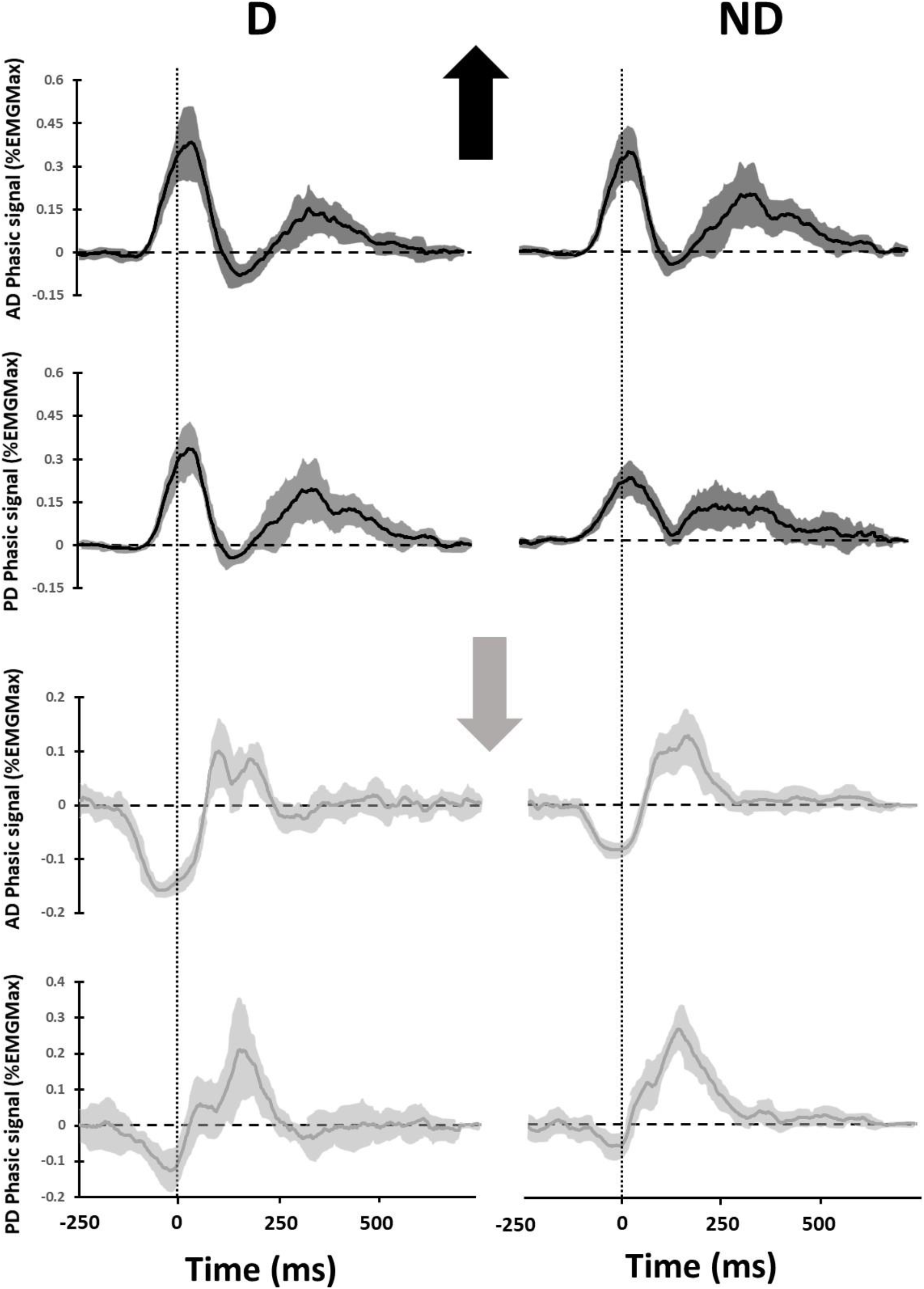
Illustration of the mean (+SD) phasic EMG patterns in a typical participant for each muscle and condition. Vertical arrows indicate movement directions. Left plots display D condition while right plots display ND condition. The dashed lines represent the kinematic onset of the movement.

#### Temporal organization of muscle patterns

To quantify the effect of *direction* and *arm* on the temporal organization of muscle commands, Fig.7 depicts mean (+SE) values of the agonist muscle activation onset (black circles), antagonist muscle activation onset (white circles), and negativity (deactivation) of the antigravity-muscle onset (grey squares). The onset of agonist muscle (F = 49.63; p < 1e-6; η_p_^2^ = 0.63), antagonist muscle (F = 417.53; p < 1e-6; η_p_^2^ = 0.94), and negativity (F =249.47; p < 1e-6; η_p_^2^ = 0.90) significantly varied with movement direction. Agonist and antagonist muscles were activated later for downward than for upward movements, independently of the arm. As previously observed (Gaveau et al. 2021), negativity onset was earlier for downward (at the beginning of the acceleration phase) than for upward movements (at the beginning of the deceleration phase), for both arms. There were no main effects of *arm* or interaction effects of *arm x direction* on agonist onset (F = 0.006; p = 0.94; η_p_^2^ = 2e-4 and F = 0.43; p = 0.52; η_p_^2^ = 0.01, respectively), antagonist onset (F = 0.002; p = 0.96; η_p_^2^ = 8.3e-5 and F = 1.32; p = 0.26; η_p_^2^ = 0.04, respectively), and on negativity onset (F = 0.50; p = 0.48; η_p_^2^ = 0.02 and F = 3.30; p = 0.08; η_p_^2^ = 0.11, respectively). Thus, the temporal organization of muscle patterns mirrors the well-known direction effect on the temporal organization of arm kinematics (Gaveau et al. 2011, 2014, 2016, 2021; Gentili et al. 2007; Hondzinski et al. 2016; Poirier et al. 2020; Le Seac’h and McIntyre 2007; Yamamoto et al. 2019, 2016; Yamamoto and Kushiro 2014). However, it does not explain the arm effect on the directional asymmetry (*arm*\**direction* interaction) that was observed in the present study on the temporal organization of the arm kinematics.

**Figure 7.**
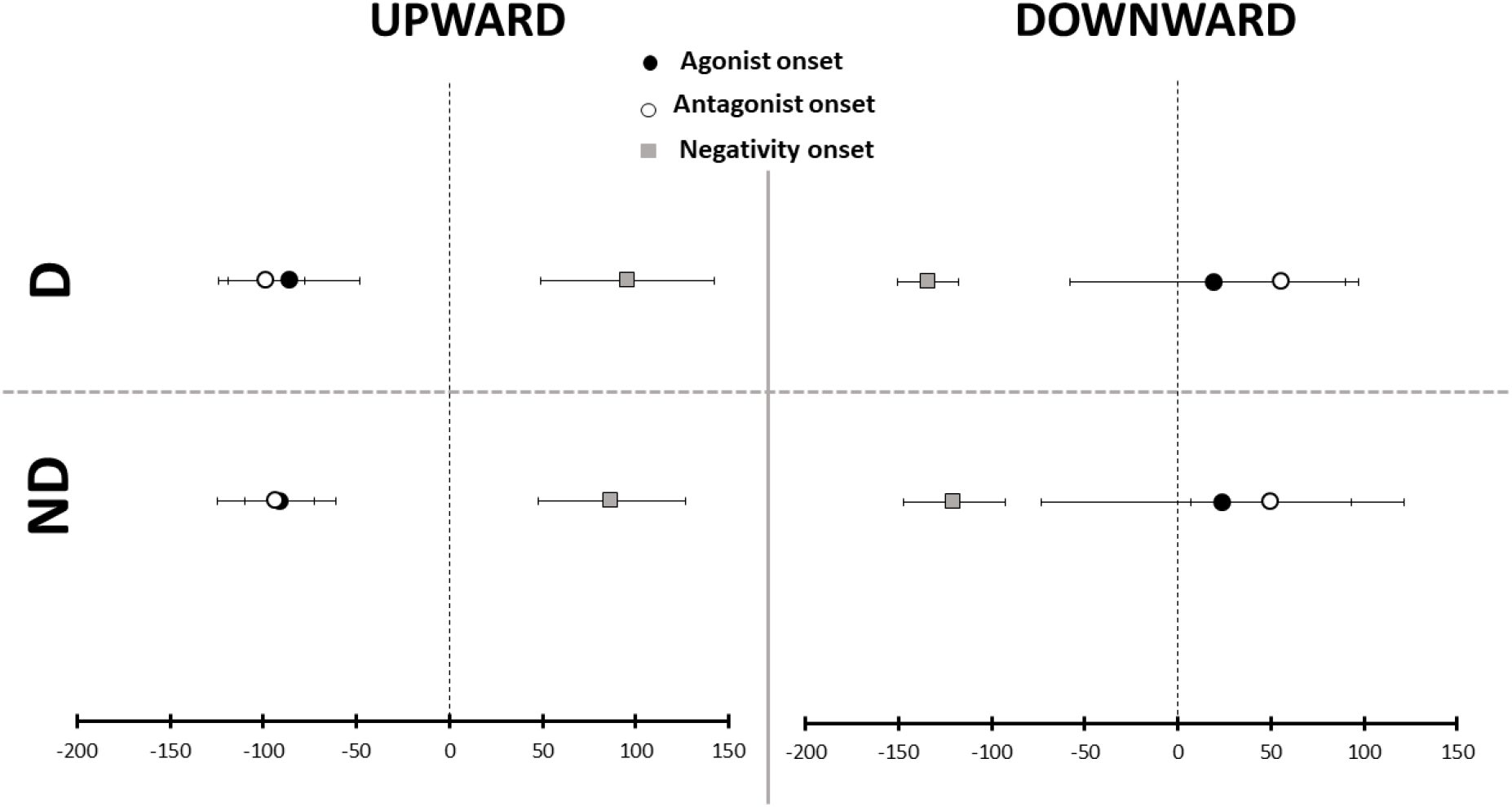
Temporal organization of muscular activation patterns. All participants’ mean (+SE) onset timing of agonist muscle (black circles), antagonist muscle (white circles), and antigravity muscle negative epoch (grey squares) are represented relative to movement’s kinematic onset (dashed lines).

#### Importance of the Negativity phenomenon

We computed the occurrence, duration, and maximal amplitude of negative epochs, for both *arms* and *directions*. Additionally, we computed a negativity index quantifying how much the antigravity-muscle was deactivated below the theoretical tonic value that would be necessary to compensate the gravity torque. These parameters are depicted in Fig.8.

**Figure 8.**
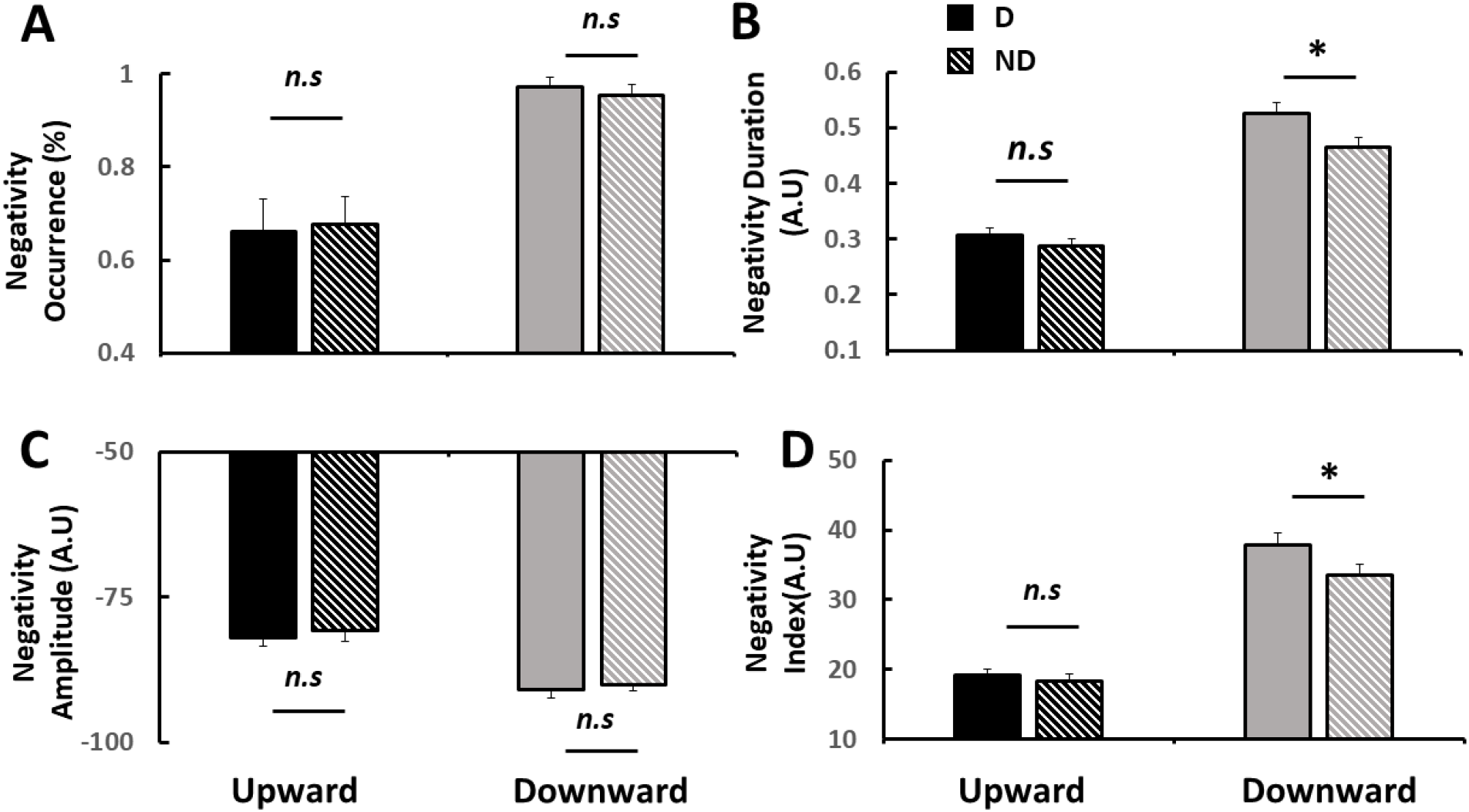
(A). Mean (+SE) negativity occurrence for upward (black) and downward (grey) movements. (B). Mean (+SE) negativity duration for upward (black) and downward (grey) movements. (C). Mean (+SE) negativity amplitude for upward (black) and downward (grey) movements. (D). Mean (+SE) negativity index for upward (black) and downward (grey) movements. For all panels, D and ND conditions are represented by respectively solid and striped bars.

We observed a significant main effect of *direction* on negativity occurrence (F = 29.61; p = 7e-6; η_p_^2^ = 0.51; Fig. 8A), negativity duration (F = 117.23; p < 1e-6; η_p_^2^ = 0.80; Fig. 8B), negativity amplitude (F = 16.50; p = 3.38e-4; η_p_^2^ = 0.36; Fig. 8C), and negativity index (F = 152.96; p < 1e-6; η_p_^2^ = 0.84; Fig. 8D). The proportion of trials with a negative epoch was 67% for upward movements and 97% for downward movements. For comparison, this proportion was 30% and 60% for upward and downward movements in gravity-muscles, respectively. The duration and the amplitude of negative epochs, as well as the negativity index, were found larger in downward compared to upward movements. *Arm* condition significantly influenced negativity duration (F = 6.91, p = 0.01, η_p_^2^ = 0.19) and index (F = 5.24; p = 0.03; η_p_^2^ = 0.15), but did not significantly influence negativity occurrence (F = 0.001; p = 0.97; η_p_^2^ = 4.4e-4), nor amplitude (F = 0.66; p = 0.42; η_p_^2^ = 0.02). Importantly, *arm x direction* interaction was found to be significant for negativity duration (F = 7.03; p = 0.01; η_p_^2^ = 0.20) and index (F = 8.57; p = 0.007; η_p_^2^ = 0.23) but did not significantly influence negativity occurrence (F = 0.24; p = 0.63; η_p_^2^ = 0.008) nor amplitude (F = 1.81; p = 0.19; η_p_^2^ = 0.06). Post-hoc comparisons revealed that negativity duration and negativity index was higher in the D than in the ND arm for downward (p = 0.006, Cohen’s d = 0.84; p = 0.004; Cohen’s d = 1.00; for duration and index, respectively) but not for upward movements (p = 0.99, Cohen’s d = 0.23; p = 0.97, Cohen’s d = 0.13; for duration and index, respectively). Similarly, it was found that, whereas negativity index didn’t differ between arms for upward movements (p = 0.97, Cohen’s d = 0.13), it was significantly higher for D compared to ND arm for downward movements (p = 0.004; Cohen’s d = 1.00; see Fig. 8D).

These results indicate that the amount of muscular deactivation – which is thought to demonstrate the use of gravity torque to minimize muscle effort – is more important with the D than with the ND arm, during downward movements only.

## Discussion

We probed a formerly hypothesized superiority of the dominant (D) over the non-dominant (ND) arm for gravity effects optimization in motor control. To this aim, we analyzed kinematic and electromyographic parameters of vertical arm movements. We observed larger directional asymmetries (upward vs downward differences) in the temporal organization of movements performed with the dominant arm, compared to the non-dominant one. Our results also revealed larger negative epochs on the phasic activity of antigravity muscles for downward movements performed with the dominant arm. Overall, the dominant arm seems to superiorly take advantage of gravity effects to minimize muscle effort.

The present results add to the existing literature investigating gravity-related arm movement planning and control (for reviews see Berret et al., 2019 and White et al., 2020). Studies investigating single-degree-of-freedom vertical arm movements, to isolate gravity effects, revealed specific features that include: i) direction-dependent kinematics (e.g., durations to peak acceleration and peak velocity are shorter for upward than for downward movements; Gentili et al., 2007; Le Seac’h and McIntyre, 2007; Crevecoeur et al., 2009; Gaveau and Papaxanthis, 2011; Gaveau et al., 2011, 2014, 2016, 2021; Yamamoto and Kushiro, 2014; Yamamoto et al., 2016, 2019; Hondzinski et al., 2016; Poirier et al., 2020) and ii) negative epochs on the phasic activity of antigravity muscles during the acceleration and deceleration of downward and upward movements respectively (Gaveau et al. 2021). Optimal control model simulations explained these features as a strategy that takes advantage of gravity effects to minimize muscle effort (Berret et al. 2008; Crevecoeur et al. 2009; Gaveau et al. 2014, 2016, 2021).

In the present study, we took advantage of this knowledge to investigate arm motor control lateralization. We observed directional asymmetries on the arm kinematics for both the dominant and the non-dominant arm (Fig.4). We also observed specific and systematic negative epochs on the phasic activations of muscles in both arms (Fig.8). These results first indicate that the effort-minimization process is functional with both the dominant and the non-dominant arm.

Although functional, we found that some markers of the effort minimization process were reduced in the non-dominant arm compared to the dominant one. First, the size of the directional asymmetry (upward vs downward) on end-point kinematics was significantly reduced in the non-dominant arm. Smaller directional asymmetries can denote lesser exploitation of gravitational torque to save muscle effort. Indeed, using a smoothness/effort composite cost function, model simulations have demonstrated that diminishing the weight of effort in the composite cost reduces directional asymmetries (Gaveau et al. 2016). It is worth mentioning that lessening the anticipated value of gravity acceleration (Gaveau et al. 2016) or the arm’s mass (Gaveau et al. 2014) would also predict decreased directional asymmetries. However, because we did not observe any arm effect (nor *arm*\**direction* interaction) on end-point errors, the interpretation of a reduced weight force anticipation seems unlikely. Second, during downward movements, the negativity index was found smaller in the non-dominant side. Theoretically, a negative epoch in phasic EMG represents a period during which the muscle is less activated than required to compensate gravity torque (D’Avella et al. 2006; Flanders and Herrmann 1992; Gaveau et al. 2021). Thus, a larger amount of negativity corresponds to a greater muscle deactivation. Since muscle activation can be considered as a proxy for muscle effort (Carrier et al. 2011; Praagman et al. 2003), the smaller negativity index observed in the non-dominant arm further suggests that the non-dominant arm motor system is less efficient at taking advantage of gravity effects to minimize muscle effort.

These results substantiate the concept of a motor control specialization within each cerebral hemisphere, where the dominant hemisphere would be specialized in open-loop processes (Bagesteiro and Sainburg 2002; Schaffer and Sainburg 2017; Yadav and Sainburg 2011, 2014). Dominant over non-dominant superiority has been demonstrated for inertial, intersegmental, and environmental dynamics predictive control (Bagesteiro and Sainburg 2002; Sainburg 2002; Sainburg and Kalakanis 2000; Yadav and Sainburg 2014). Some studies suggest that the superiority of the dominant arm for predictive control may lead to advantageous motor strategies that optimize some hidden costs, such as movement smoothness, variance, or muscle effort (Schaffer and Sainburg 2017; Yadav and Sainburg 2011). The present results provide direct experimental support for this hypothesis.

We observed differences between arms during downward but not during upward movements. Specifically, the temporal organization of arm kinematics significantly differed between arms during downward but not during upward movements. Similarly, the negativity index was significantly larger in the dominant arm for downward but not upward movements. One can speculate that this distinction is owed to the respective temporal occurrences of muscle effort savings. During downward movements, gravity effects are advantageous in the early acceleration phase, whereas they are advantageous in the late deceleration phase during upward movements. Accordingly, negative phases occur early during downward movement (before movement onset) and later during upward movements (around 100ms after movement onset, in the second half of the motion). Thus, the effort-minimization (or gravity effects optimization) signatures that were found to be reduced in the non-dominant arm happen early in the movement. They likely reflect between arms differences in feedforward control. Research in motor control has shown that optimal motor control may be implemented via predictive mechanisms, but also via a feedback control policy (Pruszynski and Scott 2012; Scott 2004; Todorov and Jordan 2002). In accordance with previous results demonstrating non-dominant side deficits for predictive control (Bagesteiro and Sainburg 2002; Sainburg 2002; Schaffer and Sainburg 2017; Yadav and Sainburg 2011, 2014), our results suggest that optimal motor control in the non-dominant arm is unaffected when arm dynamics control is mediated by feedbacks but becomes less efficient when control relies on predictive mechanisms.

The analysis of end-point errors may further support this idea. We found positive constant errors during upward movements (target overshoot) and negative constant errors during downward movements (target undershoot). These results are in accordance with those of previous works investigating end-point errors in vertical aiming (Elliott et al. 2010, 2014, 2017; Lyons et al. 2006). These studies suggested that participants often undershoot downward targets because of the important cost of time and energy associated with the reversal corrector movement that is then performed against gravity. The present results indicate that such late final error optimization is preserved during the motion of the non-dominant arm.

## References

Bagesteiro LB, Sainburg RL. Handedness: Dominant arm advantages in control of limb dynamics. J Neurophysiol 88: 2408–2421, 2002.

Berret B, Darlot C, Jean F, Pozzo T, Papaxanthis C, Gauthier JP. The inactivation principle: Mathematical solutions minimizing the absolute work and biological implications for the planning of arm movements. PLoS Comput Biol 4: e1000194, 2008.

Berret B, Delis I, Gaveau J, Jean F. Optimality and modularity in human movement: From optimal control to muscle synergies. In: Springer Tracts in Advanced Robotics. Springer, Cham, p. 105–133, 2019.

Buneo CA, Soechting JF, Flanders M. Muscle activation patterns for reaching: The representation of distance and time. J Neurophysiol 71: 1546–1558, 1994.

Carrier DR, Anders C, Schilling N. The musculoskeletal system of humans is not tuned to maximize the economy of locomotion. Proc Natl Acad Sci U S A 108: 18631–18636, 2011.

Carson RG, Goodman D, Elliott D. Asymmetries in the discrete and pseudocontinuous regulation of visually guided reaching. Brain Cogn 18: 169–191, 1992.

Crevecoeur F, Thonnard J-L, Lefèvre P. Optimal integration of gravity in trajectory planning of vertical pointing movements. J Neurophysiol 102: 786–96, 2009.

D’Avella A, Fernandez L, Portone A, Lacquaniti F. Modulation of phasic and tonic muscle synergies with reaching direction and speed. J Neurophysiol 100: 1433–1454, 2008.

D’Avella A, Portone A, Fernandez L, Lacquaniti F. Control of fast-reaching movements by muscle synergy combinations. J Neurosci 26: 7791–7810, 2006.

Elliott D, Dutoy C, Andrew M, Burkitt JJ, Grierson LEM, Lyons JL, Hayes SJ, Bennett SJ. The influence of visual feedback and prior knowledge about feedback on vertical aiming strategies. J Mot Behav 46: 433–443, 2014.

Elliott D, Hansen S, Grierson LEM, Lyons J, Bennett SJ, Hayes SJ. Goal-Directed Aiming: Two Components but Multiple Processes. psycnet.apa.org, 2010. doi:10.1037/a0020958.

Elliott D, Lyons J, Hayes SJ, Burkitt JJ, Roberts JW, Grierson LEM, Hansen S, Bennett SJ. The multiple process model of goal-directed reaching revisited. Neurosci. Biobehav. Rev.72Pergamon: 95–110, 2017.

Flanders M, Herrmann U. Two components of muscle activation: Scaling with the speed of arm movement. J Neurophysiol 67: 931–943, 1992.

Flanders M, Pellegrini JJ, Soechting JF. Spatial/temporal characteristics of a motor pattern for reaching. J Neurophysiol 71: 811–813, 1994.

Flowers K. Handedness and controlled movement. Br J Psychol 66: 39–52, 1975.

Gaveau J, Berret B, Angelaki DE, Papaxanthis C. Direction-dependent arm kinematics reveal optimal integration of gravity cues. Elife 5, 2016.

Gaveau J, Berret B, Demougeot L, Fadiga L, Pozzo T, Papaxanthis C. Energy-related optimal control accounts for gravitational load: comparing shoulder, elbow, and wrist rotations. J Neurophysiol 111: 4–16, 2014.

Gaveau J, Grospretre S, Berret B, Angelaki DE, Papaxanthis C. A cross-species neural integration of gravity for motor optimization. Sci Adv 7: eabf7800, 2021.

Gaveau J, Paizis C, Berret B, Pozzo T, Papaxanthis C. Sensorimotor adaptation of point-to-point arm movements after spaceflight: the role of internal representation of gravity force in trajectory planning. J Neurophysiol 106: 620–9, 2011.

Gaveau J, Papaxanthis C. The temporal structure of vertical arm movements. PLoS One 6: e22045, 2011.

Gentili R, Cahouet V, Papaxanthis C. Motor planning of arm movements is direction-dependent in the gravity field. Neuroscience 145: 20–32, 2007.

Hondzinski JM, Soebbing CM, French AE, Winges SA. Different damping responses explain vertical endpoint error differences between visual conditions. Exp Brain Res 234: 1575–1587, 2016.

Jayasinghe S AL, Sarlegna FR, Scheidt RA, Sainburg RL. Somatosensory deafferentation reveals lateralized roles of proprioception in feedback and adaptive feedforward control of movement and posture. Curr. Opin. Physiol. 19Elsevier: 141–147, 2021.

Lyons J, Hansen S, Hurding S, Elliott D. Optimizing rapid aiming behaviour: Movement kinematics depend on the cost of corrective modifications. Exp Brain Res 174: 95–100, 2006.

Oldfield RC. The assessment and analysis of handedness: The Edinburgh inventory. Neuropsychologia 9: 97–113, 1971.

Papaxanthis C, Pozzo T, McIntyre J. Kinematic and dynamic processes for the control of pointing movements in humans revealed by short-term exposure to microgravity. Neuroscience 135: 371–383, 2005.

Poirier G, Papaxanthis C, Mourey F, Gaveau J. Motor Planning of Vertical Arm Movements in Healthy Older Adults: Does Effort Minimization Persist With Aging? Front Aging Neurosci 12, 2020.

Praagman M, Veeger HEJ, Chadwick EKJ, Colier WNJM, Van Der Helm FCT. Muscle oxygen consumption, determined by NIRS, in relation to external force and EMG. J Biomech 36: 905–912, 2003.

Pruszynski JA, Scott SH. Optimal feedback control and the long-latency stretch response. Exp. Brain Res. 218Springer: 341–359, 2012.

Roy EA, Elliott D. Manual asymmetries in visually directed aiming. Can J Psychol 40: 109–121, 1986.

Roy EA, Kalbfleisch L, Elliott D. Kinematic analyses of manual asymmetries in visual aiming movements. Brain Cogn 24: 289–295, 1994.

Sainburg RL. Evidence for a dynamic-dominance hypothesis of handedness. Exp Brain Res 142: 241–258, 2002.

Sainburg RL. Convergent models of handedness and brain lateralization. Front Psychol 5: 1092, 2014.

Sainburg RL, Kalakanis D. Differences in control of limb dynamics during dominant and nondominant arm reaching. J Neurophysiol 83: 2661–2675, 2000.

Schaffer JE, Sainburg RL. Interlimb differences in coordination of unsupported reaching movements. Neuroscience 350: 54–64, 2017.

Scott SH. Optimal feedback control and the neural basis of volitional motor control. Nat. Rev. Neurosci. 5: 532–544, 2004.

Le Seac’h AB, McIntyre J. Multimodal reference frame for the planning of vertical arms movements. Neurosci Lett 423: 211–215, 2007.

Todorov E, Jordan MI. Optimal feedback control as a theory of motor coordination. Nat Neurosci 5: 1226–1235, 2002.

White O, Gaveau J, Bringoux L, Crevecoeur F. The gravitational imprint on sensorimotor planning and control. J Neurophysiol, 2020. doi:10.1152/jn.00381.2019.

Woytowicz EJ, Westlake KP, Whitall J, Sainburg RL. Handedness results from complementary hemispheric dominance, not global hemispheric dominance: evidence from mechanically coupled bilateral movements. J Neurophysiol 120: 729–740, 2018.

Yadav V, Sainburg RL. Motor lateralization is characterized by a serial hybrid control scheme. Neuroscience 196: 153–167, 2011.

Yadav V, Sainburg RL. Limb dominance results from asymmetries in predictive and impedance control mechanisms. PLoS One 9: e93892, 2014.

Yamamoto S, Fujii K, Zippo K, Kushiro K, Araki M. The kinetic mechanisms of vertical pointing movements. Heliyon 5: e02012, 2019.

Yamamoto S, Kushiro K. Direction-dependent differences in temporal kinematics for vertical prehension movements. Exp Brain Res 232: 703–711, 2014.

Yamamoto S, Shiraki Y, Uehara S, Kushiro K. Motor control of downward object-transport movements with precision grip by object weight. Somatosens Mot Res 33: 130–136, 2016.

